# DeepDrug: A general graph-based deep learning framework for drug-drug interactions and drug-target interactions prediction

**DOI:** 10.1101/2020.11.09.375626

**Authors:** Qijin Yin, Xusheng Cao, Rui Fan, Qiao Liu, Rui Jiang, Wanwen Zeng

## Abstract

Computational approaches for accurate prediction of drug interactions, such as drug-drug interactions (DDIs) and drug-target interactions (DTIs), are highly demanded for biochemical researchers due to the efficiency and cost-effectiveness. Despite the fact that many methods have been proposed and developed to predict DDIs and DTIs respectively, their success is still limited due to a lack of systematic evaluation of the intrinsic properties embedded in the corresponding chemical structure. In this paper, we develop a deep learning framework, named DeepDrug, to overcome the above limitation by using residual graph convolutional networks (RGCNs) and convolutional networks (CNNs) to learn the comprehensive structural and sequential representations of drugs and proteins in order to boost the DDIs and DTIs prediction accuracy. We benchmark our methods in a series of systematic experiments, including binary-class DDIs, multi-class/multi-label DDIs, binary-class DTIs classification and DTIs regression tasks using several datasets. We then demonstrate that DeepDrug outperforms state-of-the-art methods in terms of both accuracy and robustness in predicting DDIs and DTIs with multiple experimental settings. Furthermore, we visualize the structural features learned by DeepDrug RGCN module, which displays compatible and accordant patterns in chemical properties and drug categories, providing additional evidence to support the strong predictive power of DeepDrug. Ultimately, we apply DeepDrug to perform drug repositioning on the whole DrugBank database to discover the potential drug candidates against SARS-CoV-2, where 3 out of 5 top-ranked drugs are reported to be repurposed to potentially treat COVID-19. To sum up, we believe that DeepDrug is an efficient tool in accurate prediction of DDIs and DTIs and provides a promising insight in understanding the underlying mechanism of these biochemical relations. The source code of the DeepDrug can be freely downloaded from https://github.com/wanwenzeng/deepdrug.

## Introduction

The exploration for biomedical interactions between chemical compounds (drugs, molecules) and protein targets is of great significance for drug discovery^1^. It is believed that drugs interact with biological systems by binding to protein targets and affecting their downstream activity. Prediction of Drug-Target Interactions (DTIs) is thus important for identification of therapeutic targets or characteristics of drug targets. The abundant knowledge of DTIs also provides valuable insight towards understanding and uncovering higher-level information such as therapeutic mechanisms in drug repurposing^2^. For instance, Sildenafil was initially developed to treat pulmonary hypertension, but identification of its side effects allowed it to be repositioned for treating erectile dysfunction^3^. In addition, since most human diseases are complex biological processes that are resistant to the activity of a single drug^4, 5^, polypharmacy has become a promising strategy among pharmacists. Prediction and validation of Drug-Drug Interactions (DDIs) can sometimes reveal potential synergies in drug combinations to improve the therapeutic efficacy of individual drugs^6^. More importantly, negative DDIs are major causes of adverse drug reactions (ADRs)^7^, especially among the elderly who are more likely to take multiple medications^8^. The severe ADRs from critical DDIs may lead to the withdrawal of drugs from market, such as withdrawal of mibefradil and cerivastatin from the US market^9, 10^. Hence, accurate interactions prediction between drugs can not only ensure drug safety, but also can shed a light for drug repositioning or drug repurposing, which potentially can lower the overall drug development costs and enhance the drug development efficiency.

Over the past decade, the emergence of various biochemical databases, such as DrugBank^11^, TwoSides^12^, RCSB Protein Data Bank^13^ and PubChem^14^, has provided a rich resource for studying DTIs and DDIs for health professionals. However, prediction of novel or unseen biochemical interactions still remains a challenging task. In vitro experimental techniques are reliable but expensive and time-consuming. In silico computational approaches have received far more attention due to their cost-effectiveness and increasing accuracy in various drug-related prediction tasks^15, 16, 17, 18^. The state-of-the-art computational methods for interactions prediction rely on machine learning algorithms that incorporate large-scale biochemical data. Most of these efforts are based on the principle that similar drugs tend to share similar target proteins and vice versa^19^. Hence, the most popular frameworks formulate the prediction of DTIs and DDIs as classification tasks and use different forms of similarity functions as inputs^20^. Another common types of approach are to construct heterogeneous networks in the chemogenomics space to predict potential interactions using random walks^21^. The rise of machine learning methods, especially deep learning methods have promoted drug-related research tremendously in the last two decades, including the tasks for predicting DTIs and DDIs^22, 23^. For example, DeepDDI^16^ first generated a feature vector called structural similarity profile (SSP) for each drug, then calculated a combined SSPs of a pair of drugs by dimension reduction, i.e., PCA, from concatenation of two SSP of drugs. The combined SSPs were used for training DeepDDI model to perform DDI prediction. Similar to DeepDDI, NDD^15^ first calculated high-level features of drug by multiple drug similarities based on drug substructure, target, side effect, pathway and etc. Then it used a multi-layer perceptron for the interaction prediction based on curated features. DeepPurpose^24^ is a deep learning framework for DTI and DDI prediction tasks by integrating different types of neural network structure only using sequential inputs. DeepDTA^17^ used two convolutional neural networks to learn from compound SMILES and protein sequences to predict interactions. GraphDTA^25^ used graph neural networks and convolutional neural network to learn the high dimension features of drugs and targets separately and makes interaction prediction via fully connected layers.

In spite of these advances, there is still room for improvement in several aspects. First of all, the accurate prediction of unseen drug interactions depends heavily on the feature extraction technique or similarity kernel used. Since different forms of feature extraction or similarity kernel introduce varying amount of human-engineered bias, they often display different levels of predictive performance depending on the relevant settings and no single kernel outperforms others universally^26^. Similarity-based methods also have difficulty applied on large-scale datasets due to the significant computational complexity of measuring similarity matrices^27^. Network-based methods built upon topological properties of the multipartite graph suffer from the same problem depending on the complexity of the graph^28^. Deep learning based methods utilized either sequential or structural information, but none of them combined both information for specific drug/protein to comprehensively model the biological entities under the same framework.

In recent years, deep learning frameworks based on various of graph neural networks such as graph convolutional network (GCNs)^29^, graph attention networks (GATs)^30^, gated graph neural networks (GGNNs)^31^ and residual graph convolutional network (RGCNs)^32^ have demonstrated ground-breaking performance on social science, natural science, knowledge graphs and many other research areas^33, 34^. In particular, GCNs have been applied to various biochemical problems such as molecular properties prediction^35^, molecular generation, protein function prediction^36^. As pharmacological similarities are mainly originated and computed from not only sequential but also structural properties, graph representations of biochemical entities have shown capability of capturing the structural features of Euclidean ones without requiring feature engineering^37, 38^.

Based on these observations, we propose DeepDrug, a graph-based deep learning framework, to learn drug interactions such as pairwise DDIs or DTIs. We formulate the DDIs and DTIs prediction problem as classification/regression tasks. A key insight of our framework is that biochemical interactions are primarily determined by both the sequence and structure of the participating entities. Therefore, the performance of the predictive model ultimately depends on the accurate characterization of the sequential and structural information. Since chemical structure of drugs can be naturally represented as graphs with nodes and edges denoting chemical atoms and bonds, respectively. Protein structures also have natural graph representations with nodes and edges representing amino acids and biochemical interactions, respectively. Thus it is intuitive to employ a graph-based architecture for DeepDrug. The proposed model mainly differs from previous methods in the following two aspects: 1) By taking advantage of the natural graph representation of drugs and proteins, DeepDrug takes not only traditional sequence representation but also graph representations as inputs to learn sequential and structural features for drugs and proteins under the same framework; 2) DeepDrug utilizes RGCN module to capture the intrinsic structural information among atoms of a compound and residues of a protein and CNN module to obtain genomics information in sequence. Comprehensive experiments on different benchmark datasets demonstrate that DeepDrug can successfully learn DDIs and DTIs by combining structural features from RGCN module and sequential features from CNN module in different tasks such as binary classification, multi-class/multi-label classification and regression. A series of systematic experiments show that DeepDrug outperforms other state-of-the-art models and demonstrates high robustness under different experimental settings. By visualization of structural features of drugs in our study, we also demonstrate the effectiveness of the RGCN module in learning structural information and drug functional information that have reasonable consistency.

In summary, the contributions of this paper are summarized as follows:

– DeepDrug provides a unified framework based on RGCNs with edge features incorporation to extract structural information and CNNs to extract sequential information for both drugs and proteins for downstream DDIs and DTIs prediction. The novel design of DeepDrug architecture can capture comprehensive sequential and structural features.
– DeepDrug achieves state-of-the-art performance in both DDIs and DTIs prediction tasks. Through comprehensive experiments, including binary-class classification of DDIs, multi-class/multi-label classification of DDIs, binary-class classification of DTIs and regression of DTIs in different ratios of positive and negative samples, the superior performance of DeepDrug highlights the strong and robust predictive power of DeepDrug architecture.
– The interpretation of structural features learned from DeepDrug proves the key insight that biomedical structure may determine their function and drugs with similar structures tend to have similar targets.
– The results of drug repositioning for SARS-CoV-2 suggest that DeepDrug can be a useful tool for effectively predicting DDIs and DTIs and greatly facilitate the drug discovery process.

## Results

### Overview of DeepDrug

We developed a deep learning framework, DeepDrug, to predict drug interactions (e.g., DDIs and DTIs) by combining sequence profile and structural profile (Methods). For each input (drug or protein), we used sequence data as well as the partially available structure profile data as separate input branch to the DeepDrug model (Fig. 1). The input sequence was converted into a representation using one-hot encoding, and connected to several convolution layers. The partial structure profile was encoded as a graph, where we used DeepChem for converting drug SMILES strings into graph representations in the form of feature matrices (i.e. node feature matrices and edge feature matrices) and adjacency matrices and PAIRPred^39^ for extracting the protein PDB data into similar graph representations. Then the graph representation was fed to several residual graph convolution layers. The hidden features extracted from the sequence branch and structural branch were subsequently merged by concatenation. Finally, we concatenated the hidden features for the pair of inputs together, and a fully connected layer with sigmoid/softmax/none activation functions were used to get different types of output. The detailed architecture of DeepDrug is provided in Supplementary Fig. 1.

**Fig. 1.**
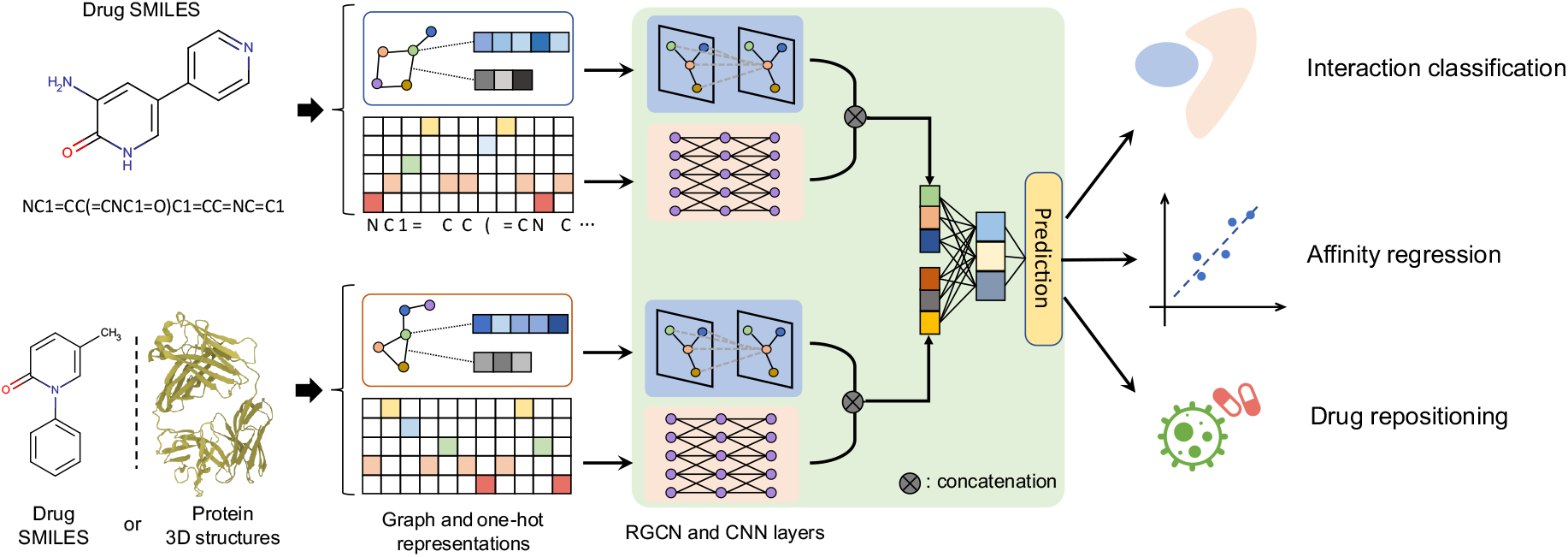
Model diagram of DeepDrug. DeepDrug is a general interaction prediction platform for DDIs and DTIs prediction. For each drug, the atom features and edge features are extracted from the SMILE sequence. For each protein in the DTI task, the node features and edge features are extracted from the corresponding pdb file (see Methods). A graph neural network module and a convolutional neural network (CNN) module are used for extracting features for drug and protein separately in the DTI task. For the DDI task, the weights in deep graph neural network are shared for a pair of drugs. The features extracted are concatenated and finally fed to a prediction module for various tasks, including interaction classification, affinity regression and drug repositioning.

### DeepDrug enables superior drug-drug interactions prediction

DDIs prediction falls into two categories: 1) binary classification where each DDI in the database was annotated as positive examples and negative examples were selected by random pairing. 2) multi-class/multi-label classification where the multi-label were obtained from annotations based on the different types of interactions (e.g., adverse effect) defined in DrugBank and Twosides. (Methods). We first evaluated the performance of DeepDrug for DDIs prediction in a binary classification setting. We benchmarked DeepDrug against six baseline methods, including random forest classification (RF) and logistic regression (LR), DeepDDI^16^, DeepPurpose^24^, NDD^15^ and AttentionDDI^40^. The baseline methods took either node feature matrix, edge feature matrix and adjacency matrix as inputs or SMILES string and similarities matrix as inputs (Methods). Four different sets of data were used for evaluation, including DDInter, DrugBank, Twosides and NDD (Methods). Our analysis showed that deep learning methods outperform similarity-based methods and traditional machine learning methods across different datasets by a large margin. Among the deep learning methods, DeepDrug consistently outperformed two other deep learning methods by achieving the highest F1 score of 0.916-0.955, highest auPRC score of 0.964-0.987 and highest auROC score of 0.971-0.988 in balanced datasets (Fig. 2A,B, Supplementary Table 1). Comparing to the second best baseline method DeepPurpose, DeepDrug achieved averaged 2.1% higher F1 score,1.3% higher auPRC score and 1.1% higher auROC score in balanced data settings.

**Fig. 2.**
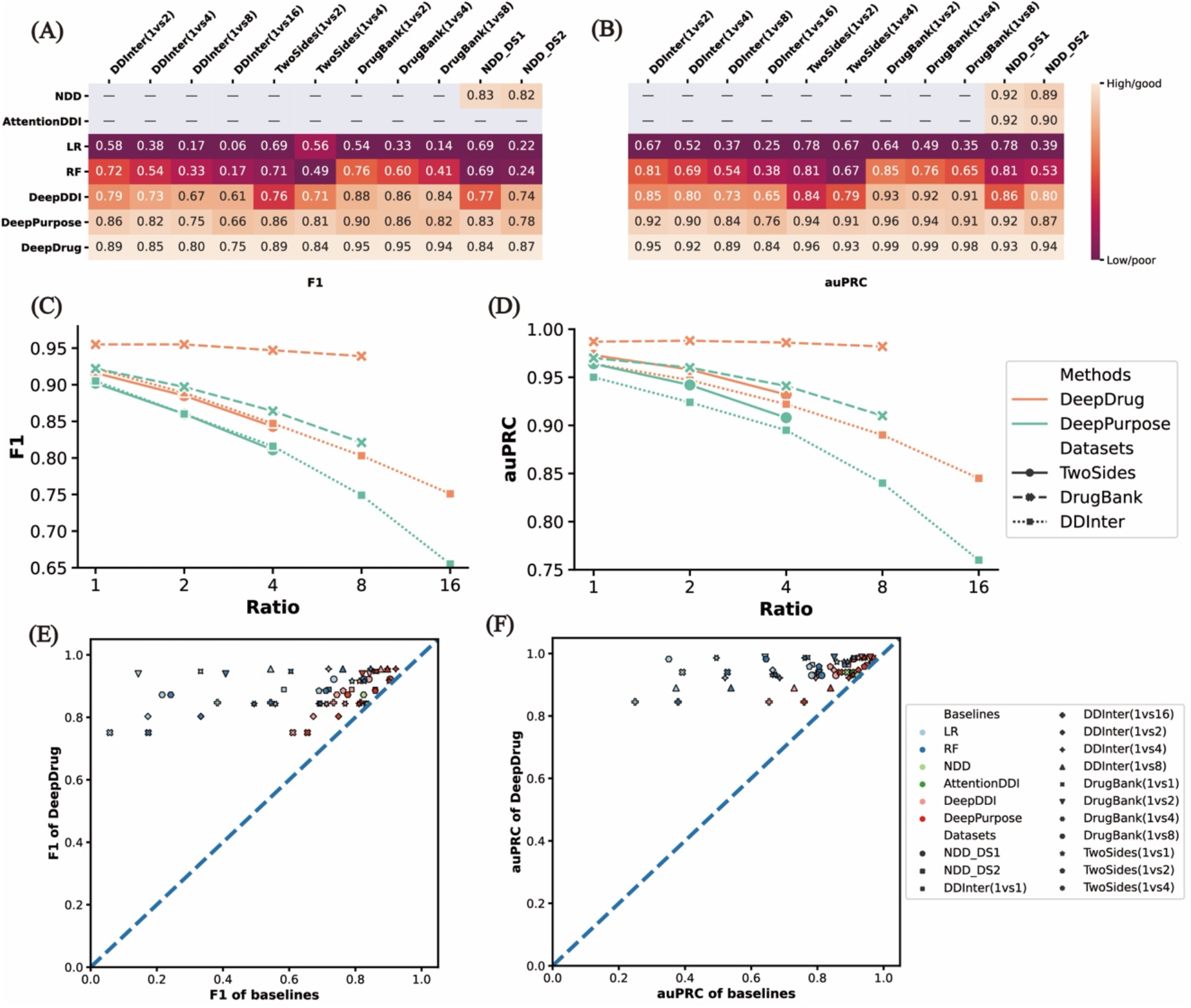
Benchmark results for DeepDrug on the binary DDI tasks. DeepDrug are benchmarked with 6 baselines on the 11 datasets in terms of F1 (A) and auPRC (B). Note that the performance scores of NDD and AttentionDDI are from the original paper and “—” indicates not applicable. The performance of DeepDrug and the best baseline method, DeepPurpose, on the unbalanced dataset are shown in (C) and (D). The x axis indicates the odds of positive to negative sample and the y axis indicates the F1 (C) and auPRC (D). Compared with DeepPurpose, DeepDrug is better to overcome the impact of unbalanced dataset. (E) DeepDrug is compared with every baseline on every dataset in terms of F1 (E) and auPRC (F). The x axis and the y axis of each dot indicate the performance of a certain baseline (indicated by dot color) and DeepDrug on a certain dataset (indicated by the dot shape).

However, Due to the rarity of occurrence of DDIs^41^, the number of known DDIs among a typical drug database is usually very low. Hence, to be more realistic and practical, we also evaluated robustness of DeepDrug with imbalanced datasets by altering the ratio between positive samples and negative samples to 1:2, 1:4, 1:8 and 1:16 based on the number of drugs in different datasets. In our case, although the auPRC scores of all comparing methods dropped, DeepDrug still outperforms other comparison methods across all datasets with different positive-to-negative ratio by achieving the highest F1 and auPRC scores (Fig. 2E-F). Specifically, DeepDrug is more robust and achieves a significantly higher performance than the second best comparison method DeepPurpose when the dataset is extremely unbalanced (Fig. 2C-D). For example, The superiority demonstrated by DeepDrug over DeepPurpose in terms of F1 score increased from 1.7% to 9.6% when the positive-to-negative sample ratio changed from 1:1 to 1:16 for DDInter dataset. To sum up, the performance of DeepDrug in terms of F1 and auPRC scores over other prediction methods demonstrated the superior ability of DeepDrug in predicting DDIs, especially with unbalanced dataset.

To further showcase the predictive capability of our model, we compare DeepDrug with other methods in multi-class/multi-label classification tasks. We conducted the classification experiments using DrugBank and Twosidess databases based on the 86 and 1317 interaction types, respectively. All of the DDI methods were evaluated using standard metrics including macro F1 score and auPRC score. In multi-class classification, DeepDrug achieved the best performance by obtaining 4.3%-5.8% higher F1 score and 4.9%-6.7% higher auPRC than the best baseline method (Supplementary Table 2). The outperformance by DeepDrug indicated the advantage of using structural representation and sequential representation of drug in DDI predictions. The same trend was observed in multi-label classification results where the introduction of 1317 types of interactions in dataset lowers F1 scores of all methods, DeepDrug demonstrated much higher F1 as 0.292 and auPRC score as 0.265 than second best method DeepPurpose (F1 score 0.227 and auPRC score 0.191) (Supplementary Table 2, Supplementary Fig. 2A,B,C). DeepDrug was shown to be superior and robust in both binary and multi-class/multi-label classification of DDIs. Therefore, unlike DeepPurpose that only used the SMILES sequence information, DeepDrug exploited both structural information from a novel graph representation and sequence information from SMILES string, which is potentially capable of learning the underlying structural properties to gain better performance.

### DeepDrug accurately identifies drug-target interactions

Although proteins generally have more intricate structures than chemical drugs due to their three-dimensional arrangement of sequence residuals, they can still be effectively represented by 3D graphs. We first classified the DrugBank DTI dataset with binary labels and benchmarked DeepDrug against six baseline methods, including RF and LR, DeepPurpose, CPI^42^, MolTrans^43^ and TransformerCPI^44^ (Methods). Similarly, the baseline methods took either node feature matrix, edge feature matrix and adjacency matrix or SMILES string as inputs (Methods). Three benchmark datasets were introduced, including BindingDB, DAVIS and KIBA (Methods). The benchmark experimental results also showed the same trend as DDIs tasks that deep learning methods dominated the DTIs prediction tasks. DeepDrug again obtained the best performance across all three deep learning methods by achieving an average auPRC of 0.811 in the above three datasets, compared to 0.788 of the second best baseline DeepPurpose (Fig.3A-C, Supplementary Table 3). Noticeably, DeepDrug and DeepPurpose were the only two deep learning methods that were applicable in the largest BindingDB dataset while the transformer-based method TransformerCPI failed due to the low computational efficiency. The superior performance of DeepDrug in DTIs prediction tasks indicated that the graph-based representation of drug can be regarded as a general framework for boosting preidcition performance in various drug-related tasks.

**Fig. 3.**
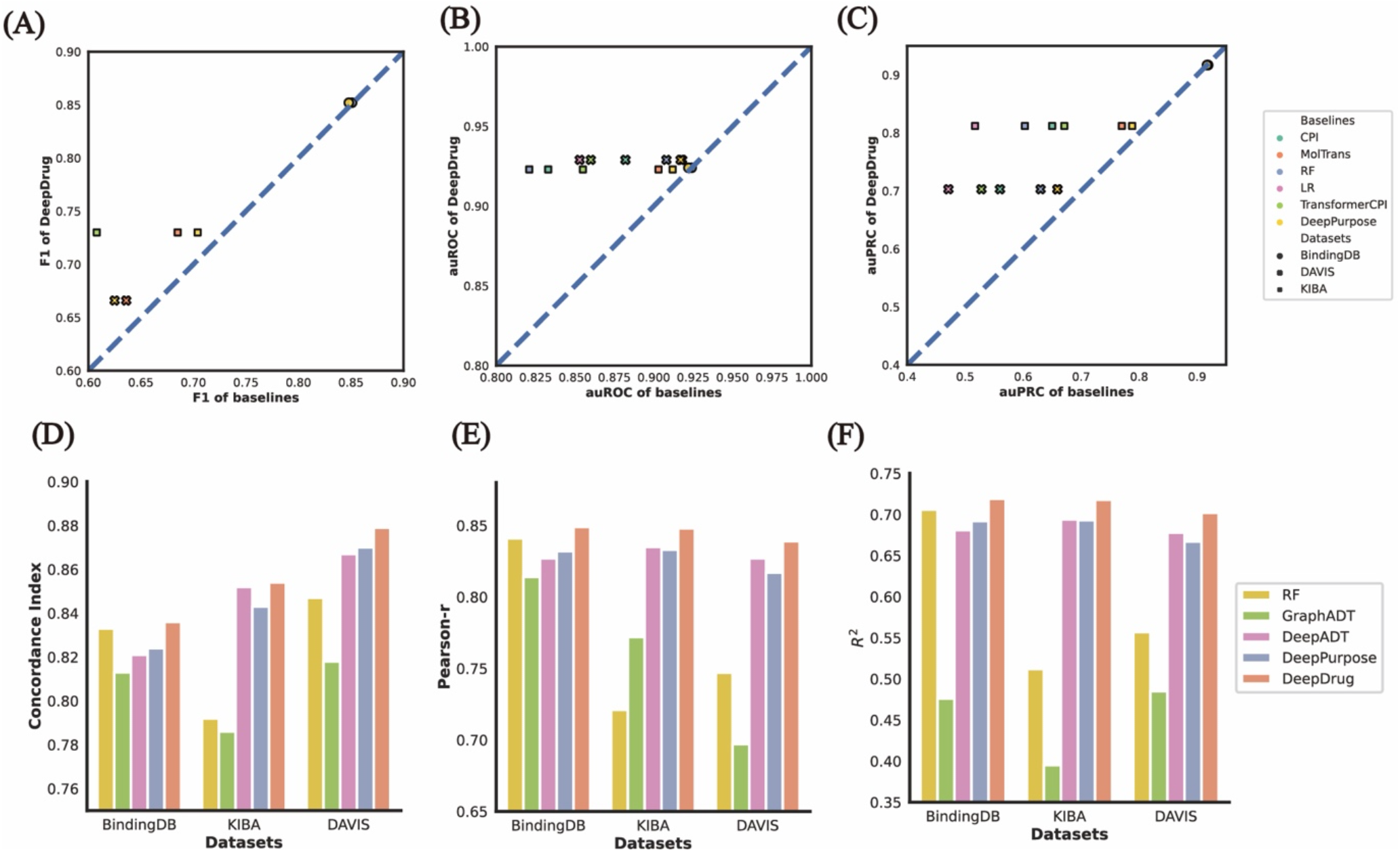
Benchmark results for DeepDrug on the DTIs and DTIs tasks. DeepDrug is benchmarked with 6 baselines on three datasets in terms of F1 (A), auROC (B) and auPRC (C) in the DTI tasks. DeepDrug is benchmarked with 3 baselines on the three datasets in terms of concordance index (D), Pearson correlation (E) and R2 (F) in the DTIs tasks.

To further illustrate the predictive capability of our model, we compared DeepDrug four baseline methods, including GraphADT, DeepADT, DeepPurpose and RF, in DTIs regression settings. We conducted the regression experiments in the same three datasets (BindingDB, DAVIS and KIBA^45^) based on the Kd value (kinase dissociation constant). All of the comparing methods were evaluated using standard metrics including concordance score, Pearson r score and R^2^ score. Again, DeepDrug achieved the best performance in terms of the three evaluation measurements compared to baseline methods (Fig. 3D-F, Supplementary Table 4). Specifically, DeepDrug achieved the highest concordance score of 0.836 in BindingDB, which is 1.2% and 2.3% higher than DeepPurpose and a graph neural network based method GraphDTA, respectively. The superiority of DeepDrug was consistently observed in DAVIS and KIBA datasets. Different from GraphDTA that only updated node features in the graph convolutional layers, DeepDrug considered both node features and edge features and updated them iteratively, thus leading to a more comprehensive representation of a drug and resulting in an incremental predictive power in the DTIs tasks. The superiority of DeepDrug indicated the benefit of combining a comprehensive structural representation and sequence representation for both drugs and proteins in DTI prediction tasks.

To further explore the ability of DeepDrug in drug repositioning, we stringently separated the drugs and proteins into training and test sets, thus curating a blind test set where the drugs and proteins in test set have no overlap with training set. This task became much more challenging as both the drugs and proteins were unseen during the training process. DeepDrug demonstrated a concordance score of 0.677 and Pearson r of 0.468 in DAVIS dataset, which outperformed DeepPurpose (concordance score of 0.605, Pearson r of 0.392) by a noticeable margin (Supplementary Table 5). Therefore, by exploiting useful structural information from graph representation of drugs and proteins, DeepDrug was shown to be consistently superior over baseline methods in both classification and regression of DTIs. We then summarized that DeepDrug provided a powerful representation of both drugs and proteins by considering both the comprehensive structural information as well as the SMILES sequence information. The superior performance of DeepDrug across various settings in DDIs and DTIs prediction tasks implicated a strong generalization ability of DeepDrug in wide drug-related applications.

### Model ablation analysis

To further support the results shown in the above sections, we conducted model ablation analysis to measure the contribution of different modules used in DeepDrug architecture (Methods). In particular, we analyzed the performance of DeepDrug with respect to the following model ablation setting: presence of RGCN module and presence of CNN module. We used the binary classification task of DDIs in multiple positive-to-negative sample ratios and DTIs regression for ablation studies. It was observed that using RGCN module alone lead to a decrease in performance with 0.5%∼2.6% lower F1 score while using only CNN module resulted in a decline of 0.2%∼1.6% in F1 score (Supplementary Table 6). Similar decrease trends were noticed in terms of R^2^ and concordance score in DTI regression tasks. To summarize, RGCN module is the most important component of DeepDrug and RGCN module and CNN module are complementing each other to further improve the predictive performance, indicating the usefulness of our designed DeepDrug architecture.

We also analyzed the robustness of DeepDrug with respect to the following hyperparameter setting: choice of feature aggregation, number of hidden units in each GCN layer, the total number of GCN layers. The performance of DeepDrug was not sensitive to different feature aggregation operations, indicating the stability of different aggregation manners (Supplementary Table 7). As the number of hidden units increased significantly (e.g. 32 and higher) in RGCN layer, both evaluation metrics started to saturate. As the number of GCN layers increased, the model became insensitive to the number of RGCN and CNN layers as well (Supplementary Table 7). To sum up, DeepDrug was insensitive to most parameter choices, illustrating the robustness of the framework.

### DeepDrug embeddings reflect drug types and drug functions

We sought to demonstrate that DeepDrug effectively captured the variability of structural information in the embeddings learned from RGCN module. We first tested our method on DrugBank dataset with extra information such as drug types or drug functions. After training, the structural embeddings from DeepDrug were projected to a two-dimensional space with t-distributed stochastic neighbor embedding (tSNE) for visualization. We found that the DeepDrug embeddings exhibited clear patterns that corresponded to the underlying drug types and drug functions (Fig. 4A, Supplementary Fig. 3). We assumed that drugs that were closer in the embedding space (e.g, within the same cluster) implied the presence of certain form of higher similarity or closer relationship. To verify this, we then quantified the effectiveness of the embeddings by various evaluation settings and made direct comparisons to the state-of-the-art method DeepPurpose (Methods). The quantitative results based on unsupervised evaluation suggested that the DeepDrug embeddings consistently outperformed DeepPurpose, with 6.91% higher averaged Drug Category Enrichment Score (DCES, Methods) while the silhouette score is similar (Fig. 4B, Supplementary Table 8). This is presumably because DeepDrug takes advantage of the extra structure information of proteins. Extensive evaluations under various folds showed that the DeepDrug embeddings consistently achieved the best performance (Fig. 4C). Furthermore, to evaluate the performance of Deepdrug applied to unseen drugs, we collected 4886 unseen drugs from DrugBank website (Methods) and the DCES was 0.575 (Supplementary Table 9), which was significantly better than the background distribution (DCES of null distribution = 0.10).

**Fig. 4.**
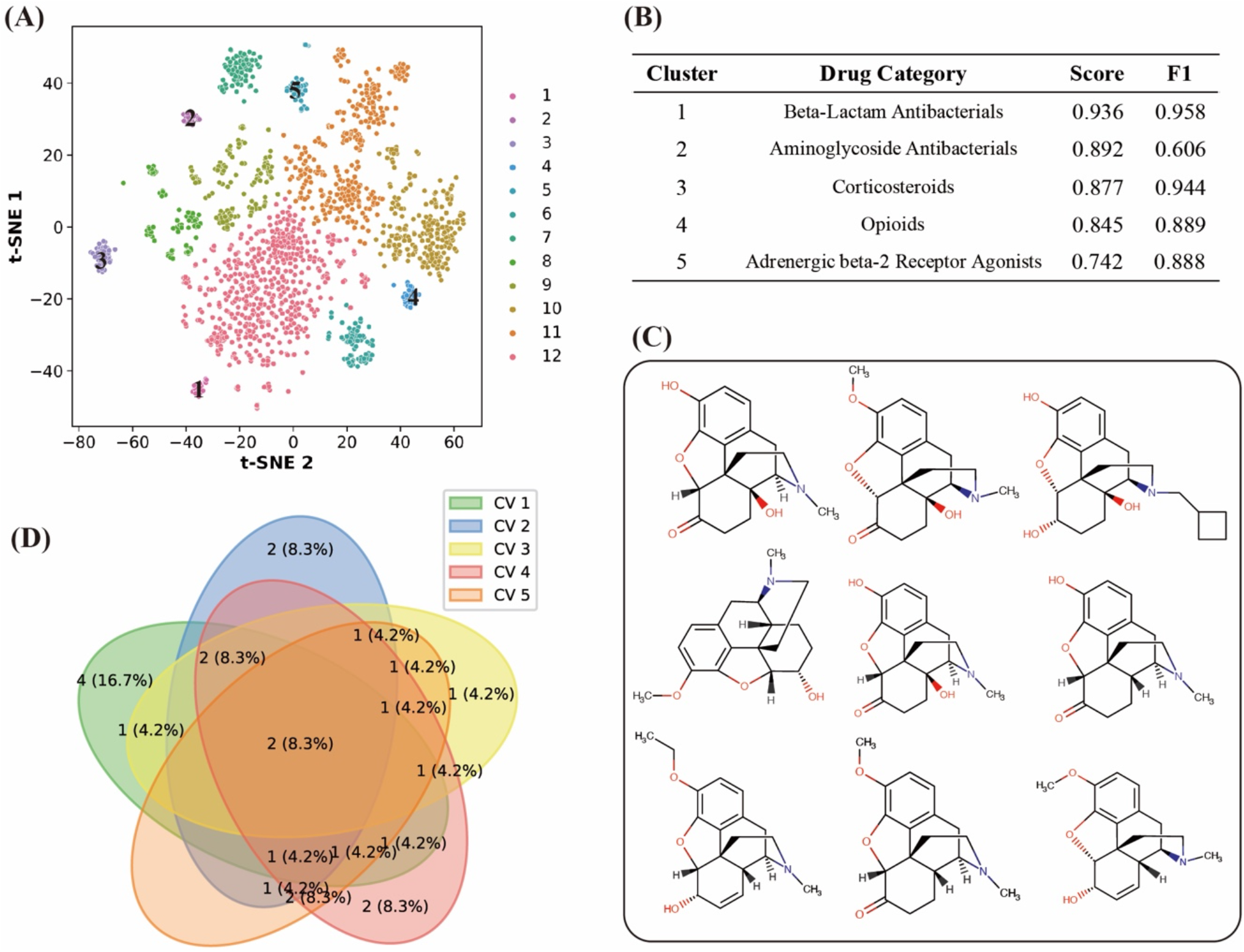
Drug embeddings for DeepDrug. (A) The t-SNE visualization for drug embeddings on the DrugBank dataset. The drug embeddings are extracted from the graph neural network module of DeepDrug. Drug embedding are then reduced dimension by t-SNE and clustered by Louvain algorithm. The enrichment of clusters with drug category are shown in (B). “Score” indicates the silhouette score of each cluster and “F1” indicates the F1-score of the cluster with best matching drug category (see Methods). Clusters are ordered by silhouette scores. The chemical structures of drugs in the cluster 4 are demonstrated in (C), indicating drug embeddings extracted by DeepDrug are topological informative. (D) Venn plot indicates the enriched drug categories of DeepDrug in each cross-validation fold are highly overlapped.

For further understanding, we isolated 12 drugs in the cluster 4 (enriched as opioids) and compared their chemical structures as well as their functionalities with other randomly sampled drugs in the dataset that were far away from the cluster. A subset of our sampled drugs are presented in Fig. 4D. The striking observation was that drugs in the cluster shared very similar structural compositions. In terms of functionality, the cluster of drugs identified by DeepDrug embeddings were highly similar among themselves as well. Out of the 28 drugs in the cluster 4, all of them were meant for pain relief (Fig. 4D). Taken together, these results demonstrated that the DeepDrug structural embeddings effectively captured the structural information which might determine the functionality of the input entities to reflect the underlying drug function. Such structural embedding capability is considered to be the main driving force to the superior performance of DeepDrug.

### DeepDrug provides therapeutic opportunities against SARS-CoV-2

To date, tens of millions of people have been infected with severe acute respiratory syndrome coronavirus 2 (SARS-CoV-2), causing the outbreak of the respiratory disease named the coronavirus disease 2019 (COVID-19). As a newly emerged member of the coronavirus family, SARS-CoV-2 is an enveloped positive-strand RNA virus, which has probably the largest genome (approximately 30 kb) among all RNA viruses. SARS-CoV-2 consists of 16 non-structural proteins (nsp1–nsp16 from ORF1a and ORF1ab), four structural proteins (spike, envelope, membrane and nucleocapsid) and nine putative accessory factors^46^. Nsp1–nsp16 forms the replicase-transcriptase complex. A primary RNA-dependent RNA polymerase Nsp12 is an essential protein that catalyzes the replication of RNA from an RNA template. The nucleocapsid (N) protein, which is mainly responsible for recognizing and wrapping viral RNA into helically symmetric structures, has been reported to boost the efficiency of transcription and replication of viral RNA, implying its vital and multifunctional roles in the life cycle of coronavirus^47^.

We next investigated whether DeepDrug was able to correctly identify the interactions of SARS-CoV-2 proteins. We constructed two drug-target positive datasets (i.e. one is expert-confirmed and one is literature-based) for SARS-CoV-2 from a recent study^46^ (Methods). In our benchmark BindingDB dataset, there were 68 SARS-CoV-2 interacting drugs and 124 proteins which were similar with these SARS-CoV-2 proteins. To obtain a stringent rule for constructing dataset, we removed those SARS-CoV-2 interacting drugs and analogous drugs from the training set that shared similar SMILE sequences (defined as drugs sequence similarities > 60%, see Methods, Supplementary Fig 4A-B), and removed proteins similar SARS-CoV-2 with protein sequence similarities > 30%. After removing these records we re-trained the DeepDrug model and combined the SARS-CoV-2 interacting drugs with the remaining 64,980 drugs to construct an independent test set. The DeepDrug prediction scores for interacting pairs and non-interacting pairs were shown in Fig 5A, we noticed that DeepDrug assigned higher prediction scores for those interacting pairs. The results showed that DeepDrug was able to distinguish expert-confirmed positive pairs from negative pairs in both of Mean and Maximum strategies (*p*-values equal to 5.06× 10^−9^ and 8.06× 10^−7^ respectively, one-side paired t-test). Results predicted on the similar templates for RCSB database, rather than the simulation structures, also show similar significant discrimination between expert-confirmed positive pairs and negative pairs (Supplementary Figure 5). In addition, the results of DeepDrug training on the original BindingDB dataset also showed similar performance (Supplementary Figure 6). Since the pairs from the negative dataset above were obtained from random pairing, there might exist potential effect drugs in the negative dataset. We observed that there were some outliers with very high affinity in the predictions of the negative pairs, which could be potential valid potential drugs. Among the top-ranked drug-protein pairs, 2 out of top-3 drugs, 5 out of top-20 drugs were already reported by literatures (Supplementary Table 10). For example, tiotropium may be effective for SARS-CoV-2 patients by reducing expressions of IL1B, IL6, IL8, RELA, NFKB1 and TNF, which is strongly activated by SARS-CoV-2 infection^48^.Such results further demonstrated the strong predictive power of DeepDrug and DeepDrug may provide therapeutic opportunities against newly found proteins such as SARS-CoV-2.

**Fig. 5.**
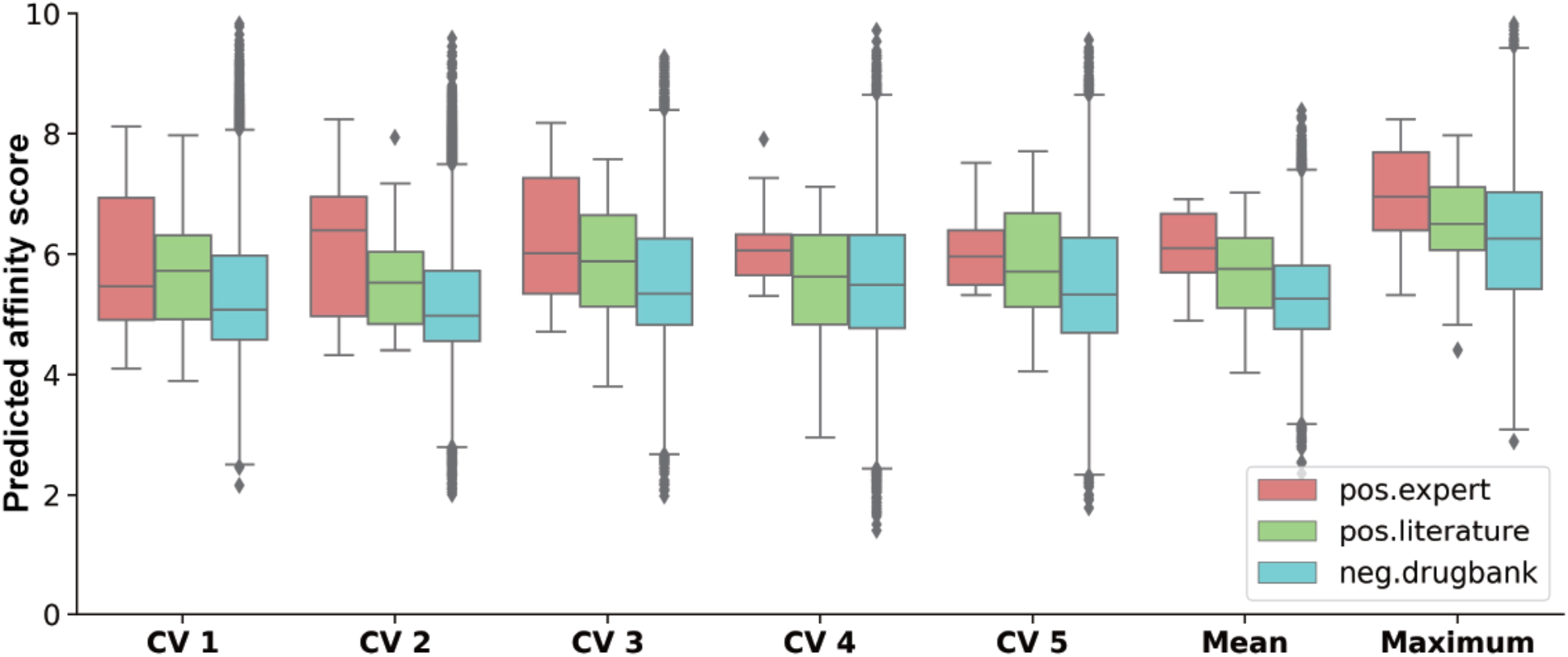
Drug repositioning for SARS-CoV-2. The performance of DeepDrug to discriminate the potential drug-SARS-CoV-2 pairs (positive dataset, i.e., “pos.expert” and “pos.literature”) and random pairs of the same drugs and SARS-CoV-2 proteins (negative dataset, i.e., “neg.drugbank”). “pos.expert” and “pos.literature” indicate two potential drug-SARS-CoV-2 pairs examined by expert checking and literature discovering individually, suggested in (Gordon, D.E., Nature 2020). The performance of DeepDrug in each cross-validation fold is shown. “Mean” indicates the final affinity scores is averaged from the 5-fold predicted affinity scores and “Maximum” indicates the final affinity scores is the maximum affinity score across all the folds. For SARS-CoV-2 proteins, simulated three-dimension structures are used.

## Discussion

In this study, we proposed DeepDrug as a novel end-to-end deep learning framework for DDIs and DTIs predictions. DeepDrug takes both topological structure information and sequence information of either drug-drug pair or drug-protein pair as inputs and utilizes RGCNs and CNNs to learn the graph representation and high-level sequence embeddings, respectively. Multi-source features are fused together to complement each other in order to achieve a superior prediction level with high accuracy. In a series of systematic experiments, DeepDrug shows superior predictive performance in different datasets under various experimental settings of DDIs and DTIs tasks. DeepDrug mainly differs from previous methods in two aspects. First, unlike most previous methods that are designed only for a specifc task such as DDIs or DTIs prediction, DeepDrug adopts a more general graph-based architecture to facilitate the prediction in both DDIs and DTIs tasks. Second, previous methods typically only used the SMILES string or drug structural information as input, DeepDrug utilizes both topological information and sequence information as inputs to form a hybrid feature representation in order to boost the prediction performance. To the best of our knowledge, DeepDrug is the first work to apply both graph convolutions and sequence convolutions to molecular representation. In addition, we demonstrated that the combination of intrinsic graph-based representation and high-level sequence embeddings are appealing for a comprehensive assessment for predicting DDIs and DTIs. Our extensive experiments highlighted the predictive power of DeepDrug and its potential translational value in drug repositioning.

We provide two future directions for improving our DeepDrug model. First, the current interaction predictions (e.g., DTIs) do not consider the causal interaction where one drug is involved in a biological or biochemical process to directly or indirectly affect a protein. Identifying such direct interactions and indirect interactions could help us better understand the related biological or chemical pathways or mecthanisms. Second, a more in-depth systematic study of the embedded features learned by RGCN module or CNN module should be carefully compared. For example, we could use correlation studies to explore the relationships between two source of features, thus providing insights in comprehensive feature interpretation.

To sum up, we introduced DeepDrug which can be served as a framework for systematically exploring the DDIs and DTIs prediction tasks with a unified model architecture. With DeepDrug, researchers could perform drug repositioning with specific target proteins. Then, one can simultaneously learn the interaction mechanism and annotate the interaction potential for every possible drug. Using large-scale pubic data, one could train an accurate and interpretable model to predict the interactions associated with human diseases (e.g., SARS-CoV-2). We hope our approach could help unveil the drug interaction mechanism and facilitate the further biochemical research.

## Method and material

### Data preparation

We collected 5 DDI benchmark datasets for evaluation. DrugBank dataset consists of 1706 drugs with 191808 drug pairs among 86 types of drug interactions. TwoSides dataset consists of 645 drugs with 63473 drug pairs among 1317 kinds of side effects. Different from the exclusive interactions of Drugbank dataset, side effects in TwoSides dataset are not exclusive, indicating that side effect prediction is a multi-label classification task. Two datasets from NDD^15^ are collected. The first one, termed NDD_DS1, is composed of 548 drugs with 300304 drug pairs, in which 97168 pairs are positive and the rest are negative. The second one, termed NDD_DS2, consists of 707 drugs with 499849 drug pairs, in which 34412 pairs are positive. DDInter^49^ dataset consists of 1493 drugs with 117608 drug pairs.

To generate a series of binary datasets from DrugBank with different positive-to-negative ratio, we considered all the pairs in the DrugBank dataset as positive samples. As for negative samples, we randomly selected drug pairs in the dataset and eliminated drug pairs that overlapped with positive samples and duplicated drug pairs. In this way, we constructed a series of binary classification datasets with positive-to-negative ratio of 1:1, 1:2, 1:4, 1:8 and 1:16.

We collected 3 DTI benchmark datasets for evaluation, including DAVIS, KIBA and BindingDB^50^ dataset. Specifically, DAVIS dataset consists of 68 drugs and 316 proteins, which constructs 21488 drug-protein pairs. KIBA dataset consists of 2111 drugs and 185 proteins, which constructs 390535 drug-protein pairs. As for BindingDB dataset, it consists of 417893 drugs and 2076 proteins, which constructs 751808 drug-protein pairs.

### Model architecture

DeepDrug was composed of RGCN modules, CNN modules, and a combined prediction module. The structural information and sequencing of each drug or protein were fed to a RGCN module and a CNN module for feature extraction, respectively. The CNN module consisted of three one-dimensional convolutional layers with 32,64 and 96 kernels respectively, followed by a one-dimensional adaptive max-pooling layer and a linear layer to extract the sequence embeddings. The kernel sizes of the convolutional layers were 4, 6 and 8 for drugs and 4,8 and 12 for proteins, respectively. Note that the adaptive max-pooling layer is aimed at reducing each kernel features across the sequence dimension by maximum function. The RGCN module was capable of learning both node embeddings and edge embeddings simultaneously by graph convolutions. Additionally, layer normalization techinique^51^ was used in node embedding layers for stabilizing the training process. The GCN module converted original node features (93 and 80 features for drug and protein respectively) to the 128-dimension features and also converted the original edge features (11 features for drug and 2 features for protein) to 128-dimension features. Borrowed from the success of deep residual network^52^, we applied convolutional residual blocks in the GCN module, which contained 22 residual blocks for DDI and 6 residual blocks for DTI. Taking the *l*-th graph convolutional residual block for an example, the node feature ***H***^*l*^ and edge feature ***E***^*l*^ were passed through layer-normalization layers,

ReLU nonlinear layers and a graph convolutional layer to obtain the processed the residual node feature 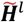 and edge feature 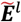, which was further added to the original input a shortcut as the final output of the graph convolutional residual block, which could be represented as 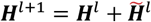 and 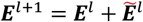. Similar to ProteinSolver graph neural network ^53^, the graph convolutional layer firstly updated edge features, i.e. 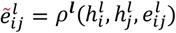 and then updated node features with updated edge features, i.e. 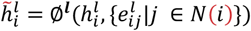. The *ρ*^***l***^, ∅^***l***^ are node and edge updating functions by aggregating the message from both node and edge in the *l*-th graph convolutional residual block. In our work, ***ρ*** was a two-layer perceptron with 256 and 128 nodes respectively, by taking the concatenation of node features and the corresponding edge features as input. The node was updated by function 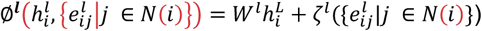, where *W*^*l*^ was learnable parameter and *ζ*^*l*^ was the message aggregation function. In our work, we chose SoftMax aggregation function ^52^.

The two RGCN or CNN modules had shared weights during DDI tasks and are independent for DTI tasks. The features extracted by these modules are concatenated together and fed to the combined prediction module, which consisted of two linear layers and a final prediction layer. The two linear layers had 128 and 32 nodes respectively, and each was followed by a batch normalization layer, a dropout layer and a ReLU nonlinear layer. The final prediction layer was a linear layer with an activation function, which was dependent on the tasks. Specifically, the Sigmoid activation function were used for binary classification task and multi-label classification task. The Softmax activation function was selected for multi-class classification task and none of activation function was used for regression task.

We used Adam optimizer with initial settings of a learning rate of 0.01, and a weight decay of 10^−4^. The dropout ratio was set to 0.1. Noticeably, we reduced learning rate by half when the loss metric on the validation dataset has stopped improving. Besides, we used Ray-project^54^ for hyper-parameters searching, including the number of the graph convolutional residual blocks, the size of channels in the graph convolutional layer (see details in Supplementary Table 11).

### Baseline methods

To evaluate the performance of DeepDrug, we benchmarked DeepDrug on multiple datasets with a 5-fold stratified cross-validation strategy for DDI tasks and DTI tasks. For classification task, we benchmarked DeepDrug with multiple baseline methods, including DeepPurpose, DeepDDI, NDD, AttentionDDI, logical regression (LR) and random forest (RF). We have modified DeepDDI slightly to make it suitable for binary classification. Note that NDD and AttentionDDI are based on multiple similarity matrices, which is not able to calculate on others dataset since the source code is not released, we have not trained and evaluated their performance on other datasets except NDD_S1 and NDD_S2. DeepPurpose is a deep learning framework for DTI and DDI prediction. We used default setting of DeepPurpose for benchmarking, i.e., “CNN” embedding for drugs and targets in DDI and DTI tasks. We have also modified DeepPurpose slightly to make it suitable for multi-class/multi-label classification tasks. For drug-target interaction task, we benchmarked DeepDrug with RF, LR, MolTrans, CPI, TransformerCPI and DeepPurpose. Note that we didn’t evaluate MolTrans, LR, TransformerCPI on BindingDB dataset due to time limitation (within 48 hours). For drug-target affinity regression tasks, we benchmarked DeepDrug with DeepDTA, GraphDTA and DeepPurpose.

### Model evaluation

F1-score, auROC (area under receiver operation curve) and auPRC (area under precision-recall curve) are used for measuring the performance in classification task. Due to the unbalance of the datasets, macro F1-score and auPRC are the more suitable metrics. For multi-label and multi-class classification, we regarded the problems as multiple binary classification tasks and calculated auROC and auPRC individually and then averaged them as the final auROC and auPRC score.

As for metrics of regression task, we used serval metrics to evaluate the performance of affinity prediction, including R^2^, Pearson correlation, and concordance index.

### Data preprocessing

We used DeepChem^55^ for drug feature calculation. Each drug is constructed with 11-dimension edge features and 93-dimension node features from SMILE strings, of which 91 features were calculated using DeepChem and the remaining two are the in-degree and out-degree of each node. As for graph feature of proteins, we firstly collected the PDB files of all proteins on the RCSB database. For each protein, we selected the longest crystal structure, i.e., the longest chain in the pdb file, as the 3D structure of the protein. Each protein is constructed with 80-dimensional node features, including amino acid features, and 2-dimension edge features (the distance of amino acids, and the angle of amino acids). Among the node features, 78 of them are calculated by PAIRPred software and the remaining two are the in-degree and out-degree of amino acids. We removed proteins without no 3D structure was found and the corresponding DTI pairs for DTI datasets.

For the DAVIS and BindingDB datasets, the binding affinity is measured by K_d_ value (kinase dissociation constant), of which the range is too large. K_d_ is transformed to logspace, i.e., pK_d_, using the formula as follows^17^:

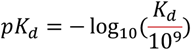

### Drug embedding interpretation

#### Drug Embedding of DrugBank DDI dataset

We firstly evaluated the drug embeddings on the DrugBank DDI dataset. For each drug in the dataset, the 128-dimension latent feature from drug feature extraction module of pretrained DeepDrug model is extracted. And then t-SNE was performed on these features for dimension reduction and Leiden algorithm^56^ is used for clustering.

The clustering results were analyzed by silhouette score and Drug Category Enrichment Score (DCES). The silhouette score is used for evaluating the clustering results. To calculate the Drug Category enrichment, 4149 drug categories were firstly collected from the DrugBank website. We noted that a drug may have multiple categories. In a certain cluster, we calculated the precision, recall and thus F1-score for each drug category. We then regarded the drug category with highest F1-score as the assigned category of this cluster. We defined the DCES as the average of the F1-scores of all clusters. The silhouette score and DCES can be used as a criterion to judge whether the drug embedding is good or not.

#### Drug Embedding of unseen drugs from DrugBank website

We collected a total of 11,172 drugs from the DrugBank website and removed drugs without any drug category, which results in 6587 drugs in total and only 1701 drugs were used for pre-training DeepDrug. Similar to above, the 128-dimension features of these 6587 drugs are extracted and dimension reduction and Leiden clustering are performed.

To calculate whether the DCES for this dataset is significant or not, we shuffled the drug categories for 1000 times and then calculated the DCES for each time, as disrupting the drug categories results in disrupting the embeddings of the drugs. We found that the mean and the standard deviation of DCES is 0.096 (+/-0.004). Thus, the DCES of DeepDrug model is significantly better than the background distribution.

### SARS-CoV-2 applications

#### Dataset

We collected two drug-target datasets for SARS-CoV-2 from a recent paper^46^, which provide a literature-based and an expert-confirmed list of drugs and target proteins for SARS-CoV-2 with 42 and 34 pairs respectively. To construct the corresponding negative samples, we randomly paired the proteins of SARS-CoV-2 and 11164 drugs from the DrugBank website. Two approaches are used for the graph features construction of SARS-Cov-2 proteins. Firstly, the simulation structures of SARS-Cov-2 proteins are provided in SARS-CoV-2 3D database^57^. Secondly, for each protein in the SARS-CoV-2, the most similar templates in the RCSB database are used as the crystal structure of the protein, which are also provided in the SARS-CoV-2 3D database^57^.

#### Drug similarity and protein similarity

To measure the similarity of two drugs, topological fingerprints of two drugs are calculated respectively by rdkit.Chem.Fingerprints.FingerprintMols.FingerprintMol function with default settings, then drug similarity is determined by rdkit.DataStructs.FingerprintSimilarity function. To measure the similarity of SARS-CoV-2 and proteins in BindingDB dataset, we first calculated the pairwise sequence similarity via EMBOSS Needle package with default setting. The drug similarity for each drug in BindingDB dataset is defined as the maximum topological similarity between this drug to each SARS-Cov-2 interacting drugs. Similar to definition of drug similarity, the protein similarity for each protein in BindingDB dataset is defined as the maximum sequence similarity between this protein to each protein in SARS-CoV-2. The drug similarity threshold is set to 0.6, which results in the removal of 64,980 drugs (15.6%). The protein similarity threshold is set to 0.3, resulting in the removal of 124 proteins (6.0%). To sum up, 128045 pairs (17.0%) are removed from BindingDB dataset to build a stringent training dataset for SARS-CoV-2 potential drug prediction.

#### Affinity prediction

We used the DeepDrug DTI models pretrained on the BindingDB dataset to predict the binding affinity of the drug-target pairs constructed above. Five-fold cross-validation pretrained model are used to make binding affinity predictions and final prediction for each pair is obtained by taking the mean and maximum values of the predictions for these 5 models.

## Code availability

DeepDrug is freely available at https://github.com/wanwenzeng/deepdrug.

## Competing interests

The authors declare no competing interests.

## Acknowledgements

We acknowledge the funding from National Key Research and Development Program of China (Nos. 2018YFC0910404 and 2020YFA0712402); National Natural Science Foundation of China (Nos. 61873141, 61721003, 62003178].

## Notes

### Competing Interest Statement

The authors have declared no competing interest.

