## Supplementary Data for "DeepDrug: A general graph-based deep learning framework for drug-drug interactions and drug-target interactions prediction"

**Supplementary Table Legends**

**Supplementary Table 1.** Benchmarking of DeepDrug on drug-drug interaction (DDI) tasks.

**Supplementary Table 2.** Benchmarking of DeepDrug on multi-class/multi-label drug-drug interaction (DDI) tasks.

**Supplementary Table 3.** Benchmarking of DeepDrug on drug-target interaction (DTI) tasks.

**Supplementary Table 4.** Benchmarking of DeepDrug on drug-target affinity regression tasks.

**Supplementary Table 5.** Benchmarking of DeepDrug on drug-target affinity regression tasks with hard cross-validation-splitting.

**Supplementary Table 6.** Model ablation on RGCN and CNN embedding channels.

**Supplementary Table 7.** Model ablation on hyperparameter, including choice of feature aggregation, number of hidden units in each GCN layer, the total number of GCN layers.

**Supplementary Table 8.** Drug embedding comparison between DeepDrug and DeepPurpose in term of Drug Category Enrichment Score (DCES) on the DrugBank dataset.

**Supplementary Table 9.** Performance of DeepDrug in term of Drug Category Enrichment Score (DCES) on the dataset curated from Drugbank website.

**Supplementary Table 10.** Predicted potential drugs for SAS-CoV-2.

**Supplementary Table 11.** Hyper-parameter searching for DeepDrug.

**Supplementary Tables**

Attached as separate csv file named Supplementary Table 1~11

**Supplementary Figure Legends**

**Supplementary Figure 1. Model detailed architecture of DeepDrug.**

**Supplementary Figure 2. Benchmark for DeepDrug on the multi-class/multi-label DDI tasks.** DeepDrug are benchmarked with 3 baselines on the 2 datasets in terms of F1 score (A), auROC (B) and auPRC (C). “TwoSides(963)” indicates the Twosides dataset with 963 types of interactions which filters out classes with less than 500 samples.

**Supplementary Figure 4. Drug embeddings for DeepDrug.** The t-SNE visualization for drug embeddings on the DrugBank dataset by DeepDrug in cross-validation (CV) fold 2 (A), CV fold 3 (B), CV fold 4 (C) and CV fold 5 (D).

**Supplementary Figure 4. Drug similarity and protein similarity.** (A) histogram of drug similarity between SAS-CoV-2 interacting drugs and drugs in the BindingDB dataset. (B) histogram of protein similarity between SAS-CoV-2 proteins and proteins in the BindingDB dataset.

**Supplementary Figure 5. Potential drug prediction for SARS-CoV-2 based on SARS-CoV-2 protein fragments.** Similar to Figure 5A, The performance of DeepDrug to discriminate the potential drug- SARS-CoV-2 pairs and random pairs of the same drugs and SARS-CoV-2 proteins. Unlike Figure 5A, the graph features for SARS-CoV-2 proteins are constructed from the similar templates in the RCSB database, rather than the simulation structures.

**Supplementary Figure 6. Potential drug prediction for SARS-CoV-2.** Similar to Figure 5A, The performance of DeepDrug to discriminate the potential drug- SARS-CoV-2 pairs and random pairs of the same drugs and SARS-CoV-2 proteins. Unlike Figure 5A, DeepDrug was trained on the original BindingDB dataset, rather than a stringent dataset.

**Supplementary Figure 1**

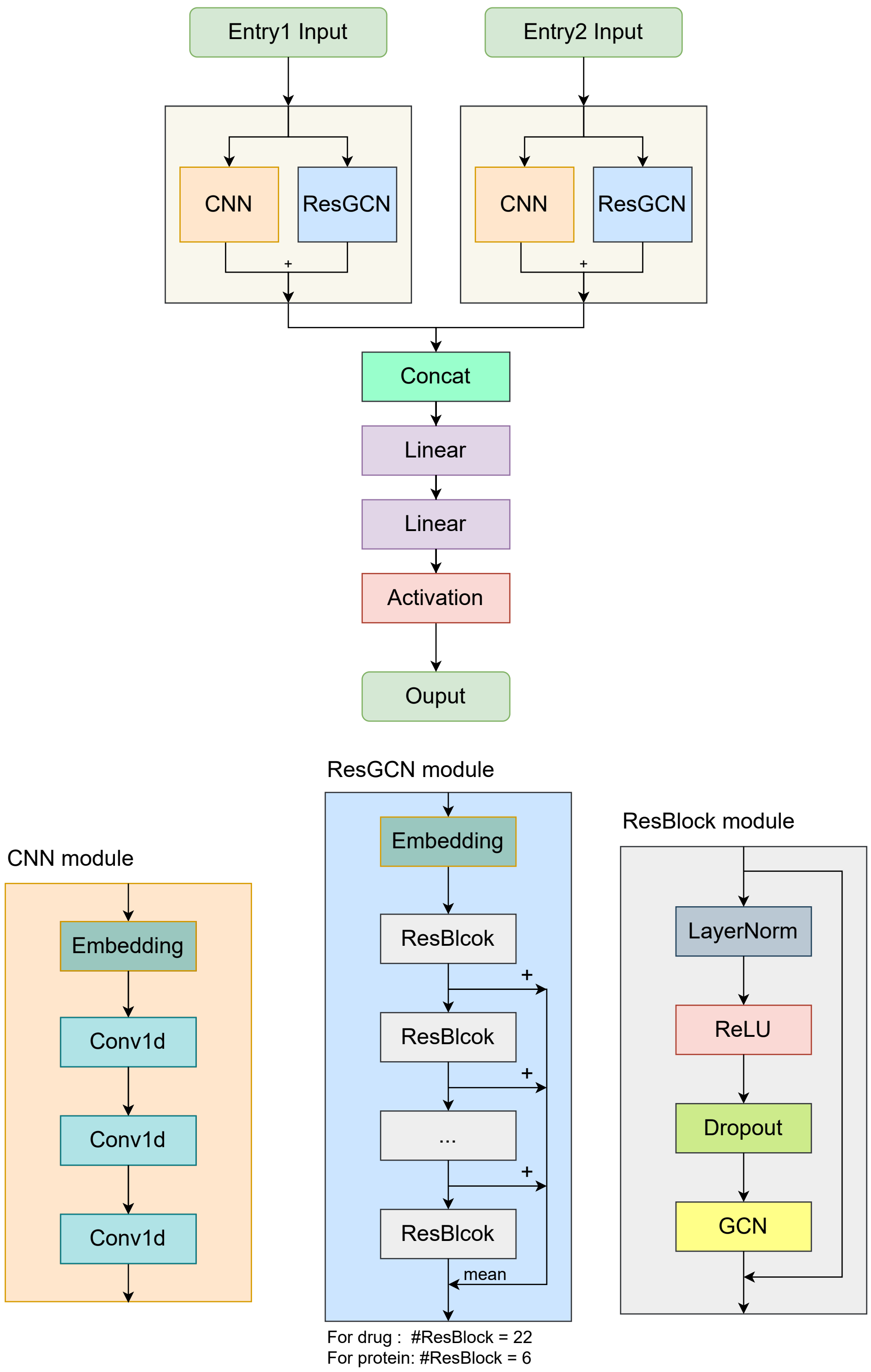

### Supplementary Figure 2

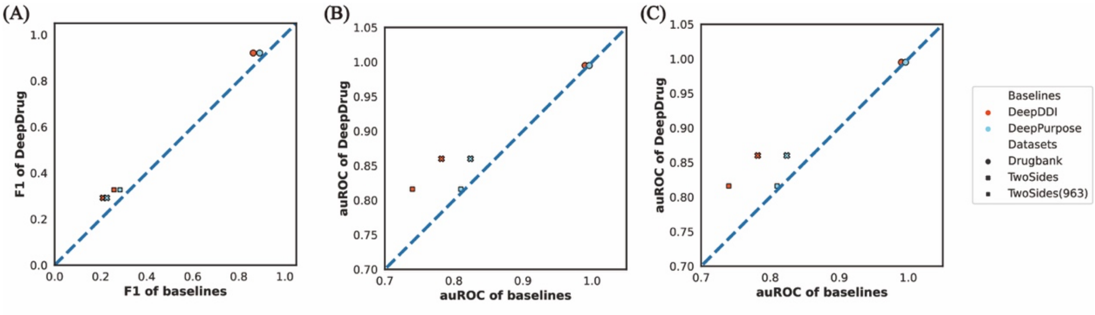

**Supplementary Figure 3**

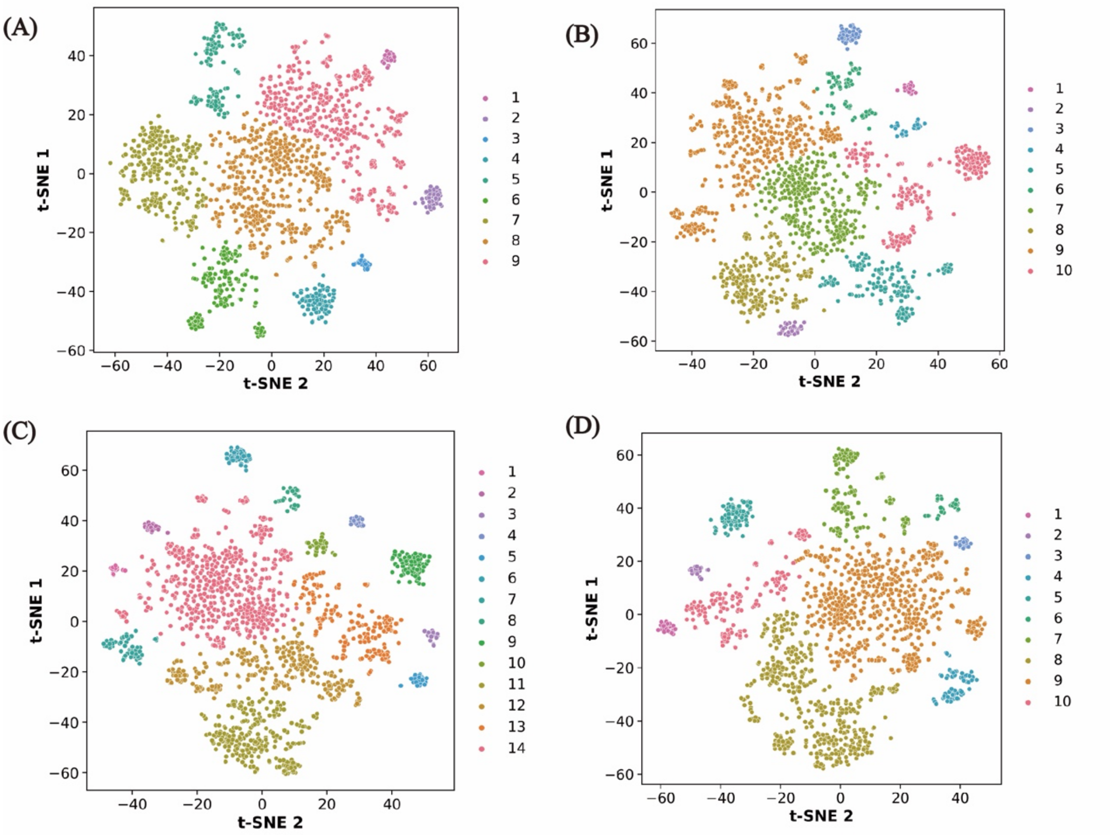

**Supplementary Figure 4**

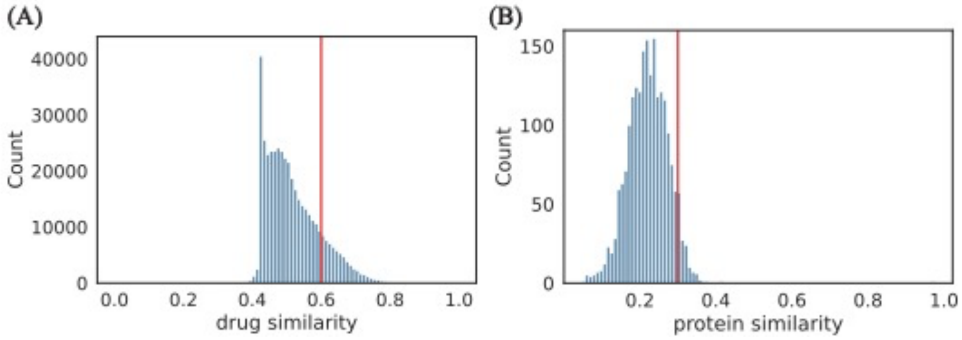

**Supplementary Figure 5**

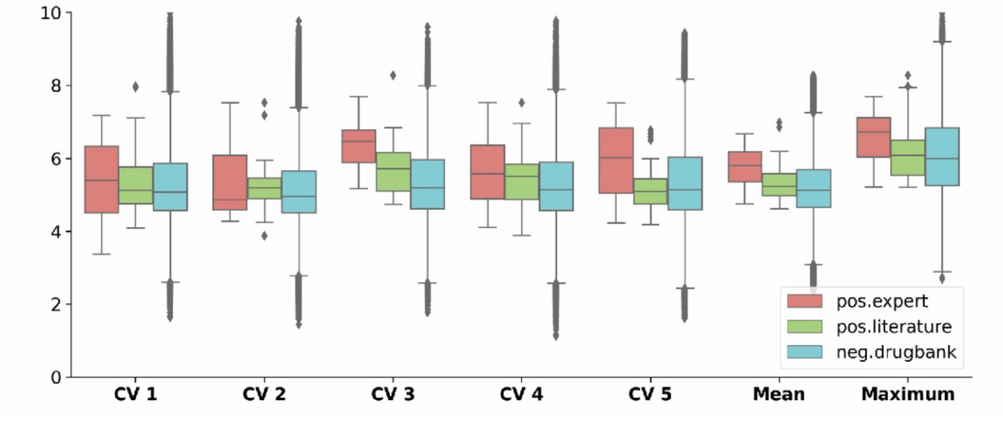

**Supplementary Figure 6**

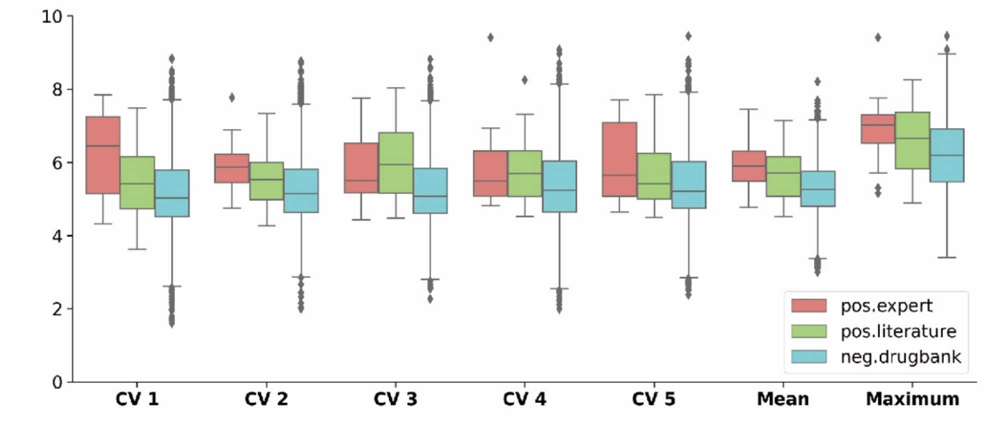
